# PERK retains a predominantly monomeric state under ER stress conditions

**DOI:** 10.64898/2026.02.04.703745

**Authors:** Konstantina Georgoula, Luo Liu, Simone Jung, Iqra Sohail, Paolo Annibale, Gabriele G. Schiattarella

## Abstract

The unfolded protein response (UPR) is a central adaptive mechanism that safeguards protein homeostasis in the endoplasmic reticulum (ER). In the heart, UPR signaling contributes to cellular remodeling and survival across a range of pathological contexts, including ischemia, pressure overload, and cardiometabolic stress. Among the three canonical UPR branches, the PKR-like ER kinase (PERK) pathway plays a critical role in modulating translational control and redox balance during stress adaptation. Despite its functional importance, the molecular dynamics of PERK activation and assembly remain incompletely understood. Here, we investigate the oligomerization behavior of PERK in living cells using advanced fluorescence microscopy. We identify a concentration-dependent mechanism of PERK self-association, as well as a distinct population of oligomeric PERK whose assembly state remains stable upon ER stress induction. These findings challenge the traditional view of stress-induced oligomerization as a prerequisite for PERK activation and suggest the existence of non-canonical modes of PERK assembly with potential regulatory significance.

## Introduction

Maintaining protein homeostasis is essential for cardiomyocyte integrity, particularly under conditions of hemodynamic, ischemic, or metabolic stress. The unfolded protein response (UPR) is a conserved signaling network that safeguards ER function by sensing and resolving disruptions in proteostasis. It is mediated by three ER-resident transducers – activating transcription factor 6 (ATF6), inositol-requiring enzyme 1 (IRE1), and protein kinase R (PKR)-like ER kinase (PERK) – each orchestrating distinct arms of the stress response. Upon accumulation of misfolded proteins and other types of stressors such as lipid stress, these sensors initiate adaptive programs that enhance protein folding capacity, degrade terminally misfolded substrates, and transiently attenuate global translation (1). In the heart, however, chronic or dysregulated UPR activation contributes to maladaptive remodeling, including cardiomyocyte dysfunction, inflammation, and fibrosis (2,3).

In the canonical model of PERK activation, ER stress triggers the dissociation of the chaperone GRP78/BiP from its luminal domain, enabling PERK dimerization and autophosphorylation. This leads to phosphorylation of eIF2α, resulting in a global reduction in protein synthesis and selective translation of stress-response factors such as ATF4 (4,5). Central to this model is the transition from inactive monomers to active dimers or oligomers as a key regulatory step. However, recent evidence challenges this view, suggesting that PERK and other UPR sensors may exist in pre-assembled states, and that ER stress does not necessarily induce major changes in oligomerization of these proteins (6,7).

To directly assess the oligomerization state of PERK in the ER membrane during UPR activation, we employed live-cell fluorescence microscopy coupled with the molecular brightness approach. This technique quantifies fluorescence intensity fluctuations to infer the stoichiometry of fluorescently tagged protein complexes at the single-cell level. Fluorescent protein-tagged PERK was used to visualize and quantify complex formation, both in isolation and in combination with other constructs, under basal and stress conditions.

## Methods

### Plasmid constructs and cloning

EYFP-PERK was created by inserting the EYFP in a pcDNA3.1 containing PERK (OHu22427, GenScript) using Gibson Assembly cloning. The EYFP sequence and the vector backbone were amplified with PCR using the following primers, with the forward primer of the vector containing a linker sequence appended to the 5’ end of the PERK gene (TCTAGA). Insert: Forward primer: 5’-AGCTCGGATCCGCCACCATGGTGAGCAAGGGCGAGGAG -3’ Insert Reverse primer: 5’-CCCGGGCTGATGGCGCGCTCTCTAGACTTGTACAGCTCGTCCATGCCGAG -3’ Vector: Forward primer: 5’-GCATGGACGAGCTGTACAAG(TCTAGA)GAGCGCGCCATCAG -3’ Vector Reverse primer: 5’-AGCTCCTCGCCCTTGCTCACCATGGTGGCGGATCC -3’ Construct BiP-mRuby was assembled to confirm the co-expression of BiP and EYFP-PERK in the cells to be imaged and analyzed. The construct was generated by inserting the sequence of BiP (Adgene, 32701) in a bicistronic vector (pVitro2, Invivogen) expressing the fluorescent protein mRuby2, using Gibson Assembly. The sequences of the primers used for amplifying the insert and the vector are the following: Insert Forward primer: 5’-CAACCGGTGATATCGCCACCATGAAGCTCTCCC -3’ Insert Reverse primer: 5’-CAAAGTGTTACCCCTCTAGACTACAACTCATCTTTTTCTGCTGTATCCT -3’ Vector Forward primer: 5’-CAGAAAAAGATGAGTTGTAGTCTAGAGGGGTAACACTTTGTACTGCG -3’ Vector Reverse primer: 5’-GCCACCAGGGAGAGCTTCATGGTGGCGATATCACCGG -3’. Constructs mTurquoise-PERK (VB230220-1087kzc), 1xEYFP-ER-RS (1xEYFP) (VB230220-1103) and 2xEYFP-ER-RS (2xEYFP) (VB230220-1110csu) were purchased and assembled from VectorBuilder. Construct mTurquoise-PERK was utilized to perform a FRET acceptor photobleaching experiment and to further validate the molecular brightness measurements regarding the oligomerization status of EYFP-PERK. Constructs 1xEYFP-ER-RS and 2xEYFP-ER-RS were required to establish molecular brightness thresholds corresponding to monomeric and dimeric protein states. Detailed sequence maps are available upon request or at the supplier’s website using the vector IDs.

### Cell culture, transfection and ER stress induction

Neonatal rat ventricular cardiomyocytes (NRVMs) were isolated and cultured in Dulbecco’s Modified Eagle’s Medium (DMEM)/M199 (3:1) (PAN-Biotech, Aidenbach, Germany) supplemented with 3% fetal bovine serum (FBS; Sigma-Aldrich, Saint Louis, MO, USA) and 100 U/mL penicillin and 100 μg/mL streptomycin (Gibco), as described previously (8). H9c2 (ATCC; CRL-1446) an HEK293 cells (Sigma-Aldrich ECACC Cat#96121229) were cultured in Dulbecco’s Modified Eagle’s Medium supplemented with 10% FBS, 2 mM L-glutamine (PAN-Biotech, Aidenbach, Germany), 100 U/mL penicillin and 100 μg/mL streptomycin. The cells were maintained at 37 °C in a humidified atmosphere with 5% CO_2_. Three days prior to imaging cells were seeded in coated, glass 8-well chambers at a seeding density of 35K cells/well (ibidi). Twenty-four hours later the plasmid constructs were transfected using Lipofectamine 2000 (ThermoFisher Scientific) according to the manufacturer’s protocol. ER stress was induced 24 h prior to imaging using thapsigargin (1 μM) or palmitate (300 μM) in H9c2 cells, thapsigargin (1 μM) or tunicamycin (1 μg/ml) in HEK cells, and thapsigargin (1 μM), tunicamycin (1 μg/ml), or phenylephrine (50 μM) in NRVMs, as indicated. Corresponding vehicle controls (0.3% DMSO or 6% BSA) were included in 300 μL of serum-free medium.

### Cell preparation for molecular brightness and FRET acceptor-photobleaching

Prior to confocal imaging for the molecular brightness approach, NRVMs, H9c2 and HEK293 cells were washed three times using DPBS (ThermoFisher Scientific) and were imaged in an imaging buffer, FluoroBrite (ThermoFisher Scientific). To ensure precise region of interest (ROI) selection without perturbations due to cell movement between frames, which may occur in live-cell imaging, HEK293 cells used in the FRET-AP experiments were fixed using 4% paraformaldehyde (PFA) for 20 min at RT, washed 3 times in DPBS and imaged in FluoroBrite.

### Microscopy

For molecular brightness experiments, cells were imaged using a LEICA Stellaris 8 confocal microscope with a white-light laser and photon counting detectors (HyD). Molecular brightness values were calculated as L=(σ^2/())-1, and normalized to the photobleached 1xEYFP-ER-RS values (1xEYFP-bl), representing a true brightness monomer, to determine the oligomeric status. Single images were collected with 512×512 pixels, a pixel size of 50 nm at a scan rate of 100 Hz. Molecular brightness values were extracted based on the spatial distribution of pixel intensities, according to the spatial brightness approach (9).

For Acceptor Photobleaching FRET, cells were imaged in a 512×512 pixel frame, 50 nm pixel size, at a scan rate of 100 Hz. FRET efficiency was measured by donor fluorescence intensities before and after photobleaching of the acceptor, calculated as E=((I_(donor after)-I_(donor before))/I_(donor after) )×100.

EYFP constructs were excited at 514 nm and fluorescence collected between 520 nm and 560 nm, mTurquoise at 440 nm and fluorescence collected between 444 nm and 523 nm, and mRuby2 at 565 nm and fluorescence collected between 580 nm and 680 nm.

### Automated ER molecular brightness analysis

The semi-automated quantification of molecular brightness is based on a previously published custom image-segmentation macro, which we adapted to segment the ER network and select membrane-only regions of interest using the Voronoi Threshold Labeler from the BioVoxxel 3D. Briefly, images were flat-field corrected, background subtracted, and contrast enhanced. ER membranes were segmented using a tubeness filter with thresholding and watershed separation. Fluorescence intensity and standard deviation were measured exclusively within these ER membrane regions, and molecular brightness of each cell was calculated using an R-based analysis pipeline (10).

### Immunoblotting

For immunoblotting, cells were lysed in ice-cold RIPA buffer (150 mM NaCl, 50 mM Tris-HCl pH 7.4, 1% Triton X-100, 0.5% sodium deoxycholate, 0.1% SDS, 5 mM EDTA) supplemented with protease and phosphatase inhibitors, according to a standard protocol. Protein samples were separated by sodium dodecyl sulfate–polyacrylamide gel electrophoresis (SDS–PAGE) on pre-cast 4–20% gradient gels (Criterion TGX, #5671095, Bio-Rad) and transferred to nitrocellulose membranes using the Trans-Blot Turbo Transfer System (#1704159, Bio-Rad). Membranes were blocked with 5% BSA and incubated with primary antibodies against PERK (C33E10, #3192, Cell Signaling Technology), ATF4 (D4B8, #11815, Cell Signaling Technology), and GAPDH (6C5, CB1001, Sigma-Aldrich), diluted in 5% BSA according to the manufacturers’ recommendations. Protein detection was performed using an Odyssey scanner (LI-COR, v3.0).

### Statistics and data analysis

Image and data analysis were performed using ImageJ, RStudio, and GraphPad Prism. Statistical analysis was performed using one-way ANOVA with Tukey’s multiple comparisons test. Differences were considered significant at p□≤ □0.05. Schematic illustrations were prepared using the freely available molecular graphics software, BioRender.

## Results and Discussion

To investigate PERK oligomerization dynamics during ER stress, we generated a PERK-specific fluorescent reporter by fusing enhanced yellow fluorescent protein (EYFP) to its luminal (N-terminal) domain. When expressed in NRVMs and H9c2 cells, EYFP–PERK localized robustly to the endoplasmic reticulum (ER), consistent with its expected subcellular distribution (**Fig. 1A, 1C**). Control constructs, including the monomeric 1xEYFP, showed comparable ER localization, supporting the specificity and proper targeting of the PERK reporter (**Supplementary Fig. 1A-C**).

**Figure 1.**
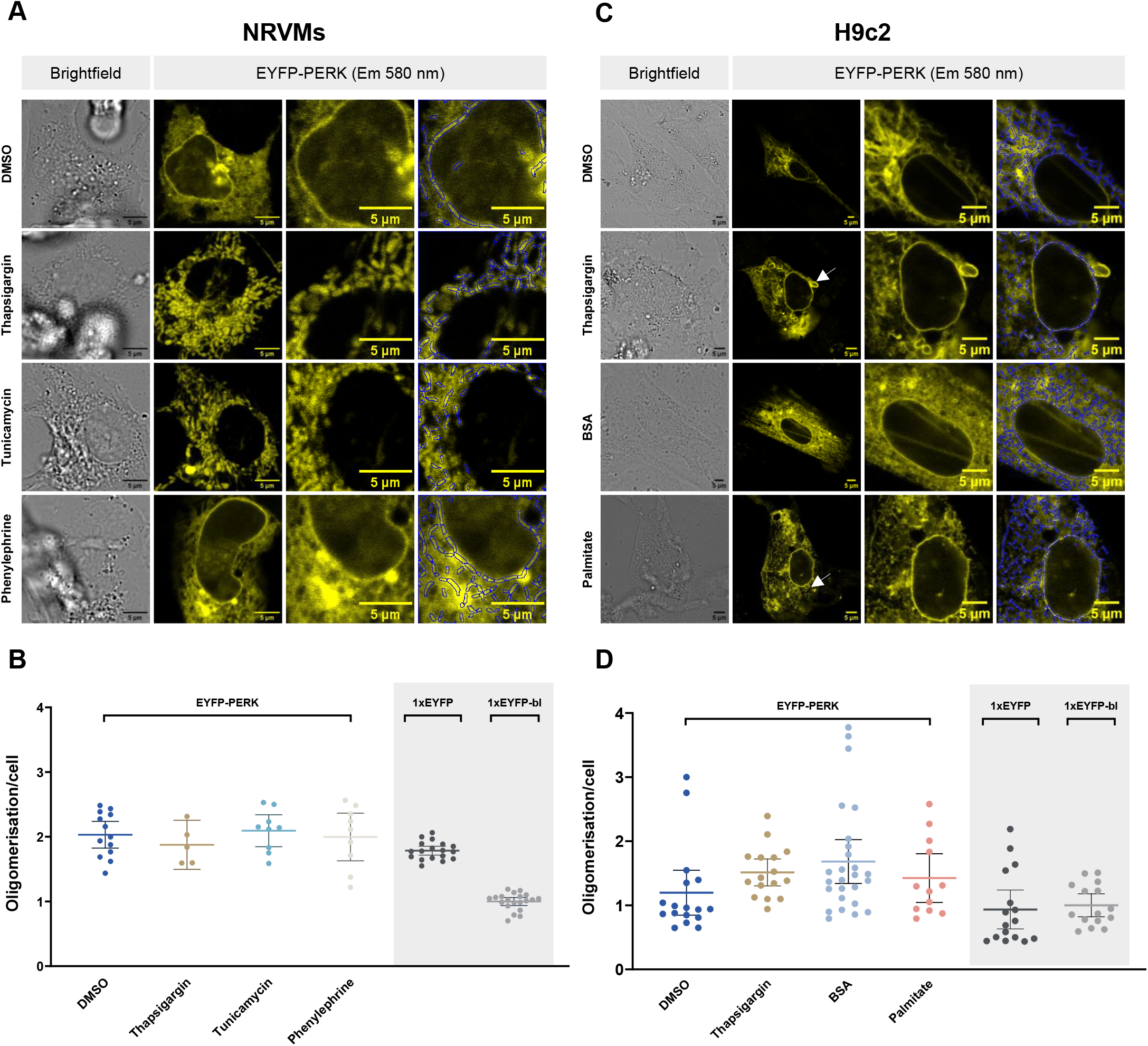
Oligomerization of PERK in response to unfolded protein response in NRVMs and H9c2 cells. **A:** Confocal microscopy images of NRVM single cells expressing the fluorescent chimaera EYFP-PERK, and how its spatial arrangement is altered by the indicated treatments. ROIs in blue highlight how areas of the Endoplasmic Reticulum are selected for subsequent analysis. **B:** Molecular Brightness analysis of EYFP-PERK together with selected monomeric control (1xEYFP) expressed in NRVMs, upon indicated treatments. The oligomerization is determined by normalizing Molecular Brightness values to the monomeric control after photobleaching (1xEYFP-bl). Data are shown as mean with 95% CI. Data points represent single cells, obtained from two independent NRVM isolation preparations, each derived from a variable number of neonatal rat hearts from approximately 25 pups. **C:** Confocal microscopy images of H9c2 single cells expressing the fluorescent chimaera EYFP-PERK, and how its spatial arrangement is altered by the indicated treatments. ROIs in blue highlight how areas of the Endoplasmic Reticulum are selected for subsequent analysis. **D:** Molecular Brightness analysis of EYFP-PERK together with selected monomeric control (1xEYFP) expressed in H9c2 cells, upon indicated treatments. The oligomerization is determined by normalizing Molecular Brightness values to the monomeric control after photobleaching (1xEYFP-bl). Data are shown as mean with 95% CI from n = 3 independent transfections per condition. Each data point corresponds to a measurement of a single cell under the indicated conditions. Statistical significance was defined as p ≤ 0.05.

To assess whether EYFP tagging, or PERK overexpression interfered with PERK function, we examined its downstream signaling activity in HEK293 cells. Although transient transfection resulted in higher EYFP–PERK levels relative to endogenous protein, ATF4 expression remained unchanged (**Supplementary Fig. 1D**), indicating that neither the fluorescent tag nor overexpression substantially altered PERK-dependent signaling.

To induce ER stress and assess its impact on PERK oligomerization we treated NRVMs with 3 mechanistically distinct ER stressors: thapsigargin, which depletes ER calcium by inhibiting the sarco/endoplasmic reticulum Ca^2+^ -ATPase (SERCA) (11), tunicamycin which induces ER stress by inhibiting N-linked glycosylation and causes misfolded protein accumulation, and phenylephrine which induces ER stress indirectly by increasing protein synthesis load on the ER (12,13) (**Fig. 1A**). In addition, H9c2 cells expressing EYFP-PERK were treated with thapsigargine or the unsaturated fatty acid palmitate, which perturbs lipid homeostasis and promotes ER expansion and protein misfolding (14). Treatments were carried out for 24 hours, with DMSO and bovine serum albumin (BSA) serving as respective vehicle controls. Confocal microscopy revealed pronounced ER remodeling in response to all stressors, characterized by the disruption of the typical tubular network and the appearance of ring-like structures (**Fig. 1A, 1C, white arrows)**. These morphological features, absent in control conditions, are consistent with sustained UPR activation and align with previous reports of stress-induced ER reorganization (15).

To quantify PERK oligomerization under stress conditions, we applied molecular brightness analysis, selectively restricted to in-plane ER membrane regions using automated segmentation and region selection tools. This spatially resolved approach minimized interference from out-of-focus signal and enabled reliable detection of subtle changes in protein assembly during UPR induction (16).

As shown in **Figure 1B and 1D**, EYFP–PERK exhibited a profile in line with that of the monomeric control (1xEYFP) at baseline, consistent with the canonical activation model. Notably, 24-hour treatment with the chosen ER stressors did not induce a significant shift in PERK oligomerization in neither NRVMs nor H9c2 cells (**Fig. 1B, 1D**). In particular, we observed no evidence for a transition toward a fully dimeric state, as instead proposed in conventional models (17). A minor increase in brightness was detected under stress, but remained within baseline variability and did not reach statistical significance.

Because oligomerization is inherently concentration-dependent and shaped by protomer affinity – governed by the microscopic association and dissociation rate constants (k_on_ and k_off_) – we next examined whether expression level affects PERK oligomerization at baseline and during ER stress. For this reason, we turned to HEK293 cells which showed markedly higher EYFP–PERK expression upon transient transfection compared to NRVMs and H9c2 cells (**Supplementary Fig. 2A**). We observed that increased expression indeed promotes higher-order assembly across both conditions (**Supplementary Fig. 2B**) and that regardless of the saturation effect we see no difference in the oligomerization mechanism of EYFP-PERK upon ER stress (**Supplementary Fig. 2D**). Notably, also the 1xEYFP control displays an increased oligomerization as expression levels increase (**Supplementary Fig. 2C**), albeit the 1:2 ration between 1xEYFP and 2xEYFP was maintained across all experiments and concentration ranges (**Fig. 2B**). In HEK293 cells, EYFP–PERK again localized appropriately to the ER (**Fig. 2A**), and displayed increased average brightness relative to the monomeric control, indicating enhanced basal oligomerization (**Fig. 2B**). However, even under these elevated expression conditions, treatment with thapsigargin and tunicamycin did not induce a measurable change in PERK oligomerization (**Fig. 2B**).

**Figure 2.**
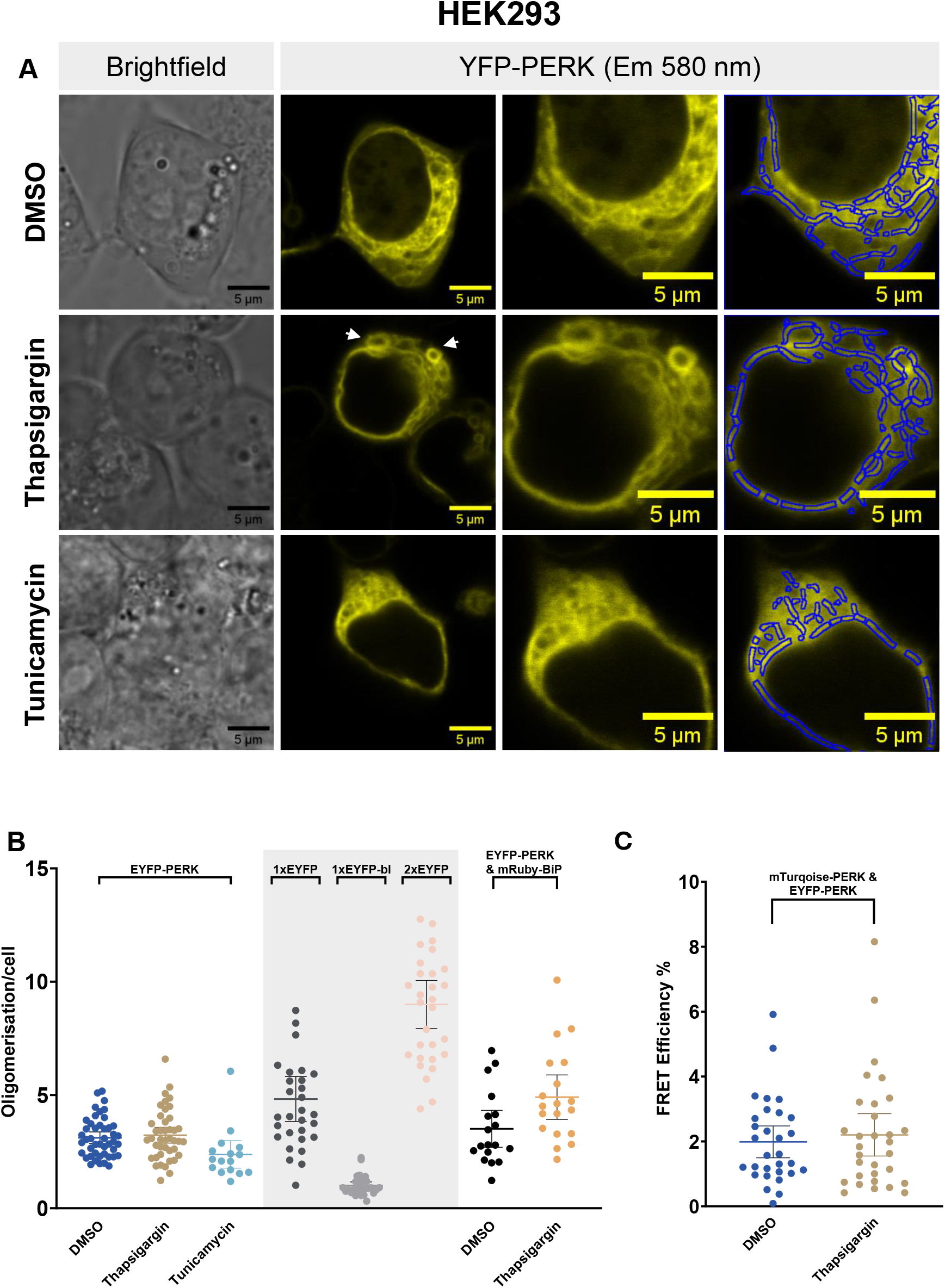
Oligomerization of PERK in response to unfolded protein response in HEK293. **A:** Confocal microscopy images of HEK293 single cells expressing the fluorescent chimaera EYFP-PERK, and how its spatial arrangement is altered by the indicated treatments. ROIs in blue highlight how areas of the Endoplasmic Reticulum are selected for subsequent analysis. **B:** Molecular Brightness analysis of EYFP-PERK together with selected monomeric control (1xEYFP), as well as dimeric control (2xEYFP), when expressed in HEK293 cells, upon indicated treatments. Molecular Brightness values are normalized to the monomeric control after photobleaching (1xEYFP-bl). **C:** Oligomeric state of PERK probed by FRET readouts in HEK293 cells. Efficiency of FRET between mTurquoise2-PERK and EYFP-PERK is monitored based on donor unquenching upon acceptor photobleaching. Data are shown as mean with 95% CI from n = 3 independent transfections per condition. Each data point corresponds to a measurement of a single cell under the indicated conditions. Statistical significance was defined as p ≤ 0.05.

Co-expression of BiP – a chaperone that binds PERK to prevent its oligomerization under basal conditions and releases it during ER stress, allowing oligomerization-led to a modest, non-significant increase in PERK oligomerization (**Fig. 2B**), consistent with a model in which BiP binding constrains PERK assembly in resting cells (5). Together, these findings indicate that PERK oligomerization is concentration-dependent – evident from its higher oligomeric state in HEK293 cells, where it is more abundantly expressed – but remains largely unresponsive to acute ER stress. Neither thapsigargin nor tunicamycin induced a measurable shift in its baseline oligomeric profile. Notably, the brightness signal of PERK was consistently lower than that of the monomeric control used in this and previous studies (7), further highlighting the influence of membrane crowding and local protein concentration on measured oligomeric states.

To independently validate our molecular brightness results, we performed Förster resonance energy transfer (FRET) analysis to assess PERK oligomerization during ER stress. PERK was tagged with a well-characterized donor–acceptor pair, mTurquoise2 and EYFP, and acceptor photobleaching experiments were conducted under baseline conditions and following thapsigargin treatment (**Fig. 2C**). Across conditions, FRET efficiency remained stable at approximately 2%, indicating no appreciable change in proximity between PERK molecules. Given that FRET detects interactions within a 1-10 nm range, these data further support the conclusion that PERK does not undergo major rearrangements in oligomeric state upon UPR activation.

In this study, we examined the oligomerization dynamics of PERK during ER stress using live-cell fluorescence microscopy – an approach previously applied only to the UPR sensor IRE1α (7). By combining molecular brightness analysis and FRET, we employed two orthogonal and quantitative techniques to monitor the oligomeric state of fluorescently labeled PERK directly in its native ER membrane environment. This live-cell approach allowed us to bypass the constraints of traditional biochemical methods and probe PERK assembly with spatial and temporal resolution under physiologically relevant conditions.

The canonical model of PERK activation posits that ER stress leads to BiP dissociation, exposing a luminal dimerization interface that facilitates PERK autophosphorylation and downstream signaling (5,18). This concept, largely derived from static assays such as immunoprecipitation, gel filtration, and crystallography, has remained widely accepted across cell types and contexts. However, our data indicate that PERK maintains a predominantly monomeric state even upon activation of the unfolded protein response, with no substantial shift toward dimerization or higher-order complex formation following treatment with established ER stressors.

Taken together, these findings support a revised view of PERK activation, one that departs from the classical model of stress-driven monomer-to-dimer transition. Our data suggest the existence of alternative, possibly pre-assembled oligomeric states that persist regardless of UPR induction. Although this finding is novel for PERK, it parallels recent work on IRE1α, where single-molecule tracking revealed the presence of inactive oligomers at baseline (7), challenging long-held assumptions about UPR sensor activation. We also show that PERK oligomerization scales with expression level – a concentration-dependent behavior consistent with other ER-resident proteins previously considered monomeric (7).

## Conclusion

This work is not confined to cell biology, but informs broader questions of stress adaptation and disease. Chronic ER stress and maladaptive UPR signaling are increasingly recognized as central features of cardiometabolic disease and heart failure with preserved ejection fraction (HFpEF), particularly in the context of metabolic overload and systemic inflammation. The notion that PERK activation can occur without large-scale oligomeric remodeling challenges existing assumptions about its regulation in stressed cardiomyocytes and opens new avenues to explore how UPR signaling contributes to the pathophysiology of HFpEF. A deeper understanding of these dynamics may help redefine how we interpret UPR activation in the failing heart – and more broadly, how we target PERK and related pathways in cardiometabolic disease.

In conclusion, this study refines the current model of PERK activation by challenging the long-held assumption that ER stress induces a shift from monomeric to dimeric or higher-order oligomeric states. Instead, our data demonstrate that PERK oligomerization is primarily shaped by expression level and local concentration, rather than by acute stress-induced remodeling. These findings extend recent revisions to the understanding of UPR sensor regulation and underscore the need to decouple protein abundance from functional activation in mechanistic studies. In the context of cardiometabolic disease and HFpEF – where ER stress and maladaptive proteostasis are increasingly implicated – these insights call for a more nuanced framework to interpret UPR signaling dynamics. Clarifying how PERK operates under physiologic and pathologic load may ultimately guide the development of more selective interventions that modulate stress adaptation without disrupting basal homeostasis.

## Funding

This work was supported in part by the following grants: DZHK (German Centre for Cardiovascular Research – 81X3100210; 81X2100282); the Deutsche Forschungsgemeinschaft (DFG, German Research Foundation – SFB-1470–A02); the European Research Council – ERC StG 101078307; HI-TAC (Helmholtz Institute for Translational AngioCardiScience) to G.G.S.; and the DFG – SFB-1470–A01 to P.A.

## Author Contributions

Konstantina Georgoula: Conceptualization, Data curation, Formal analysis, Investigation, Methodology, Validation, Visualization, Roles/Writing - original draft, Writing - review & editing

Luo Liu: Investigation

Iqra Sohail: Investigation

Simone Jung: Investigation

Paolo Annibale: Conceptualization, Funding acquisition, Methodology, Project administration, Resources, Software, Supervision, Validation, Visualization, Roles/Writing - original draft, Writing - review & editing

Gabriele Schiattarella: Conceptualization, Funding acquisition, Project administration, Resources, Software, Supervision, Validation, Visualization, Roles/Writing - original draft, Writing - review & editing

## Acknowledgements

We thank the Advanced Light Microscopy Facility of the Max Delbrück Center for Molecular Medicine. We are grateful to Richard Edel for fruitful discussion.

## Notes

### Competing Interest Statement

The authors have declared no competing interest.

