## Supplementary material for "PERK retains a predominantly monomeric state under ER stress conditions": Supplemnetary

### Supplementary Figure 1

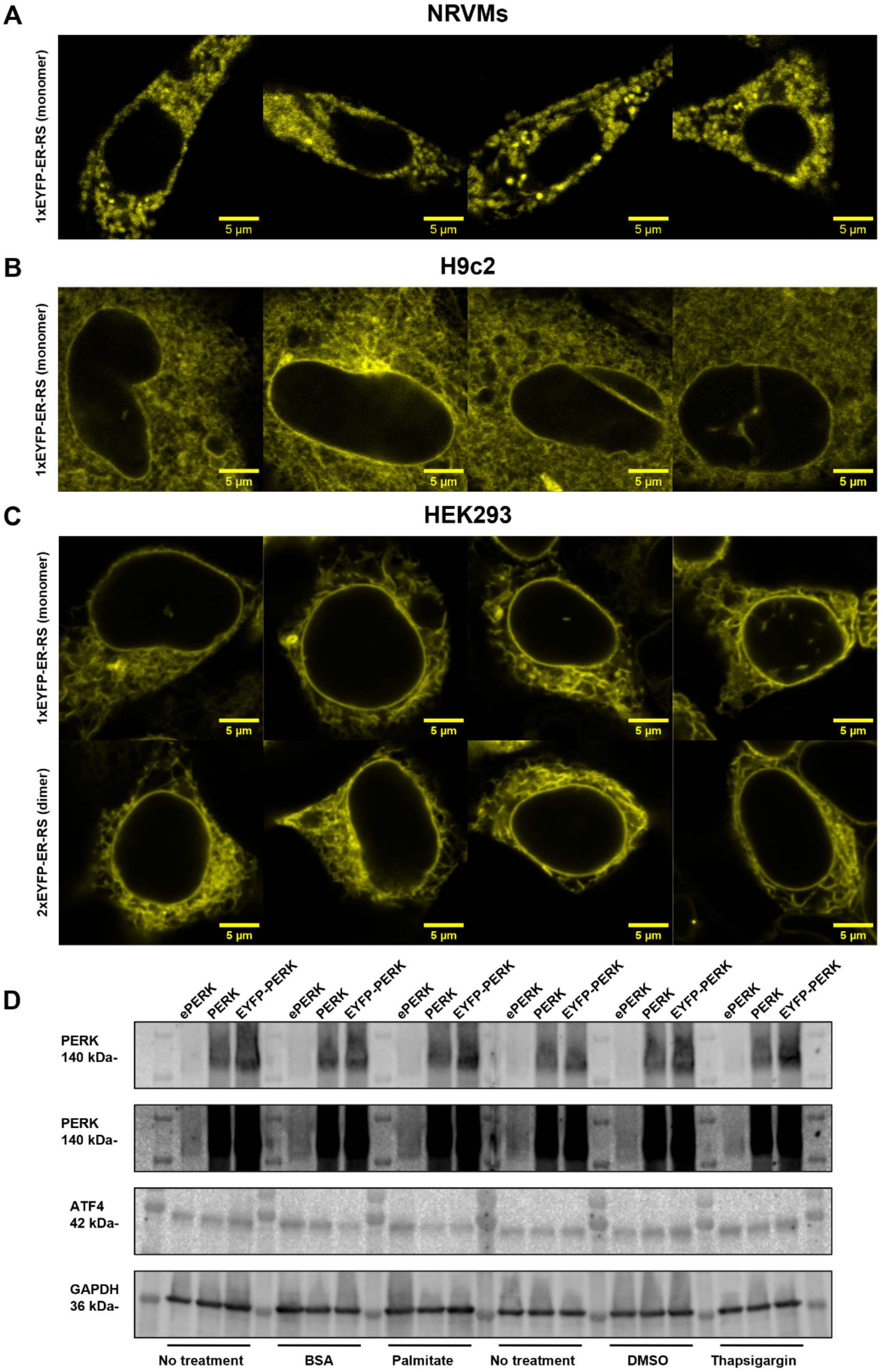


**Supplementary Fig. 1.** Expression and activation analysis of PERK variants and controls in NRVMs, H9c2 and HEK293 cells. **A:** Confocal microscopy images showing ER-localized expression of the monomeric control constructs in NRVMs. **B:** Confocal microscopy images showing ER-localized expression of the monomeric control constructs in H9c2 cells. **C:** Confocal microscopy images showing ER-localized expression of the monomeric and dimeric control constructs in HEK293 cells. **D:** Western blot analysis of PERK, ATF4, and GAPDH from HEK293 cell lysates under three expression conditions: endogenous PERK (ePERK), transfection with untagged PERK (PERK), and transfection with EYFP-tagged PERK (EYFP-PERK). Cells were either untreated or treated with BSA, palmitate, DMSO, or thapsigargin for 24 h.

### Supplementary Figure 2

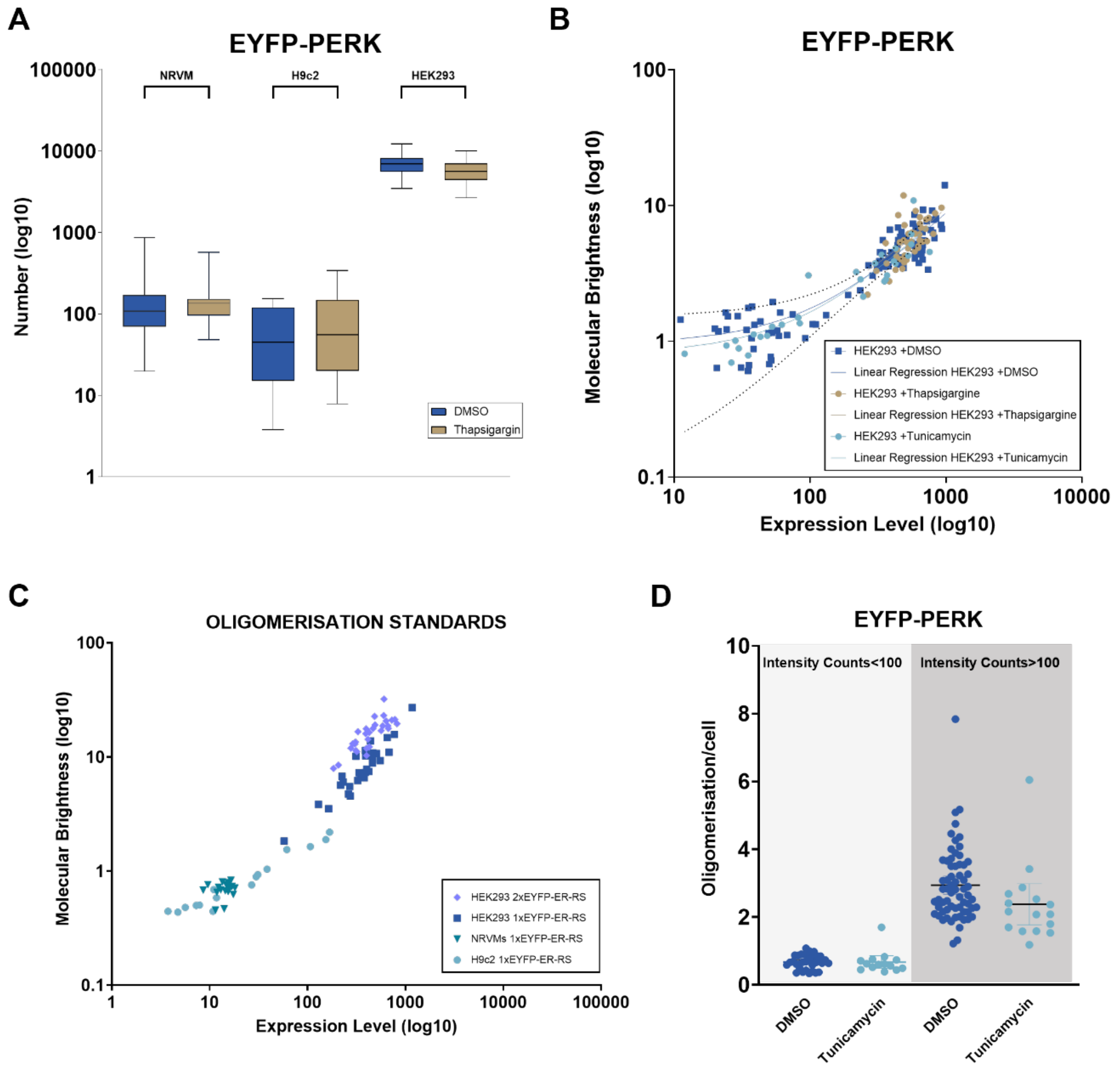


**Supplementary Fig. 2. A:** Comparison of EYFP–PERK particle numbers measured in NRVMs, H9c2, and HEK293 cells. Values are plotted on a $\log_{10}$ scale (y-axis) to account for the wide dynamic range. **B:** Scatter plot showing the relationship between molecular brightness (y-axis, $\log_{10}$ scale) and expression level (x-axis, $\log_{10}$ scale) of EYFP-PERK in HEK293 in baseline and upon ER stress induction. **C:** Scatter plot showing the relationship between molecular brightness (y-axis, $\log_{10}$ scale) and expression level (x-axis, $\log_{10}$ scale) of the monomeric control (1xEYFP-ER-RS) in NRVMs, H9c2, and HEK293 cells, and the dimeric control (2xEYFP-ER-RS) in HEK293 cells. **D:** Molecular Brightness analysis of EYFP-PERK when expressed in HEK293 cells in baseline and upon treatment with tunicamycin. Data has been grouped based on the expression level (Intensity counts<100 (light grey) and Intensity Counts >100 (dark grey). Molecular Brightness values are normalized to the monomeric control after photobleaching (1xEYFP-bl). Data are shown as mean with 95% CI from n = 3 independent transfections per condition. Each data point corresponds to a measurement of a single cell under the indicated conditions.
